# Cryptography in the DNA of living cells enabled by multi-site base editing

**DOI:** 10.1101/2023.11.15.567131

**Authors:** Verena Volf, Simon Zhang, Karen M. Song, Sharon Qian, Fei Chen, George M. Church

## Abstract

Although DNA is increasingly being adopted as a generalizable medium for information storage and transfer, reliable methods for ensuring information security remain to be addressed. In this study, we developed and validated a cryptographic encoding scheme, Genomic Sequence Encryption (GSE), to address the challenge of information confidentiality and integrity in biological substrates. GSE enables genomic information encoding that is readable only with a cryptographic key. We show that GSE can be used for cell signatures that enable the recipient of a cell line to authenticate its origin and validate if the cell line has been modified in the interim. We implement GSE through multi-site base editing and encode information through editing across >100 genomic sites in mammalian cells. We further present an enrichment step to obtain individual stem cells with more than two dozen edits across a single genome with minimal screening. This capability can be used to introduce encrypted signatures in living animals. As an encryption scheme, GSE is falsification-proof and enables secure information transfer in biological substrates.

## INTRODUCTION

DNA is the primary information storage medium of life. Its durability, replicability, and efficiency make it an ideal medium for other types of (non-biological) information. Indeed, efforts to encode information in DNA *in vitro*^1,2^ and in living systems^3–8^ have already been successful. For example, installing DNA barcodes in microbes and attaching these cells to an object, guards against DNA degradation while simultaneously allowing tagging of an object with a unique DNA barcode for object authentication^9^. The ability of DNA to propagate information over several generations is yet another desirable feature that can be leveraged in the context of information storage and transfer. Despite the many desirable features that DNA offers as a medium for information storage, propagation, and retrieval, mechanisms for ensuring information security are lacking. With genetically modified organisms becoming increasingly abundant, there is a growing demand for methods to authenticate biological substrates. Such methods would strongly benefit from secure information encoding, enabling falsification-proof genomic signatures that can be used to verify the identity of biological organisms/cell lines by a recipient and to address ownership infringements. The ability to detect whether a cell line has further been modified would be another desirable feature of such methods, akin to digital signature verification.

Motivated to address this need, we conceptualized an encryption scheme—Genomic Sequence Encryption (GSE)—that uses DNA cryptography for secure information encoding. Cryptography ensures information security by requiring a cryptographic key to decrypt the stored information. In GSE, the key comprises a list of genomic coordinates across which point mutations can be installed. Using base editors—precise and programmable genome editing tools^10,11,12,13^—we encode information by introducing multiple single-nucleotide edits across the genome. GSE provides cryptographic security through the difficulty of detecting point mutations in large genomes. With the key, information can be easily retrieved through targeted sequencing. Without the key, one would have to manually search for edits across the genome, a process prone to errors^14,15^. As an encryption scheme that is based solely on the properties of DNA sequencing and analysis, the protection GSE offers is independent of computational hardness. It is therefore not threatened by increases in computing performance, nor by the emergence of quantum computers, realizing Shor’s algorithm^16^. To the best of our knowledge, GSE is the first DNA sequencing-based encryption scheme. Prior work on information encoding in DNA relies on security through obscurity in which a short, synthesized DNA sequence is hidden in DNA ^17^. However, the system is immediately broken if the hidden information is discovered a single time or the security mechanism is otherwise revealed. Once the DNA is sequenced, short strings of sequence that do not map to a reference genome can be easily detected. Instead, here we assume DNA sequencing is abundant but rely on asymmetric sequencing cost due to encryption, analogous to asymmetric computational costs in cryptographic hashes. GSE adheres to Shannon’s Maxim, which requires a cryptographic system to remain secure, even if everything about the scheme is known (except for the key)^18^.

To implement GSE in mammalian cells, we developed a multi-site base editing protocol. We introduced targeted edits using adenine base editors (ABEs) and cytosine base editors (CBEs) to mediate the conversion of A•T→G•C and C•G→T•A, respectively. In contrast to traditional CRISPR/Cas9 editing, base editors do not rely on the induction of double-strand breaks (DSBs) and thus mitigate associated cell death and genotoxicity, as well as the insertions/deletions (indels) and chromosomal translocations that frequently occur when targeting more than one site simultaneously^13,19–21^. However, obtaining reliable editing over a high number of sites remains challenging. While precise, simultaneous modification of multiple sites on a single genome has essential applications for fundamental biology (e.g. for studying genetic interactions or long-range regulation of gene expression) as well as for biotechnology (e.g. for metabolic engineering), methods to introduce multiple edits remain laborious: Generating cells with multiple edits requires sequential editing cycles and isolating edited cells for subsequent editing rounds or extensive screening of edited clones^22^. To date, efforts to address this limitation have focused on using multi-gRNA arrays, frequently coupled with antibiotic selection^23,24^. However, the repetitive nature of guide RNAs (gRNAs) makes cloning of these constructs prone to recombination. Installing and enriching for multiple edits has proven particularly challenging in primary and stem cells, which have been of interest due to their physiological relevance and the ability to create organoid models with a specified set of mutations. Edits in these cells have been limited to no more than five sites, and efficient editing enrichment has been restricted by reliance on phenotypic selection for each site^25,26^.

In the present study, we conceptualized and implemented the cryptography scheme GSE in the genome of mammalian cells. We developed an editing protocol that enables robust, single-step, multi-site base editing using pooled editing with the capacity to encode information via simultaneous editing across >100 genomic sites. Additionally, we present an enrichment protocol that enables >25 edits in individual stem cells (*within* a single genome) with minimal screening requirements. As embryonic stem cells are commonly used for zygote injection, this step paves the way toward GSE in living animals. We demonstrate that GSE provides cryptographic security and outline an application for encrypted cell signatures that enable authentication of biological organisms. Finally, we demonstrated that GSE allows us to detect whether a cell strain has been modified in the interim by analyzing shifts in editing frequencies.

## RESULTS

We conceptualized GSE, an information encoding scheme that provides cryptographic security through the difficulty of detecting point mutations. In GSE, information is encoded across a set of genomic coordinates that represent the cryptographic key. Information is secured through the asymmetric difficulty of detecting edits with and without the key. When the key is known, targeted sequencing can be efficiently performed at high coverage to recover if the site has been edited. However, if the coordinates are unknown, all bases of the genome need to be analyzed for mutations. To implement GSE, our first goal was to achieve robust multi-site genomic editing (multiple precise edits across distinct genomic loci) in mammalian cell lines. We opted to use base editors which can be easily programmed towards multiple genomic targets in a DSB-independent manner^13,21^. Additionally, we aimed to use plasmids that each encode a single gRNA so that we could easily combine different subsets of gRNAs for information encoding (**Fig. 1a**).

**Figure 1.**
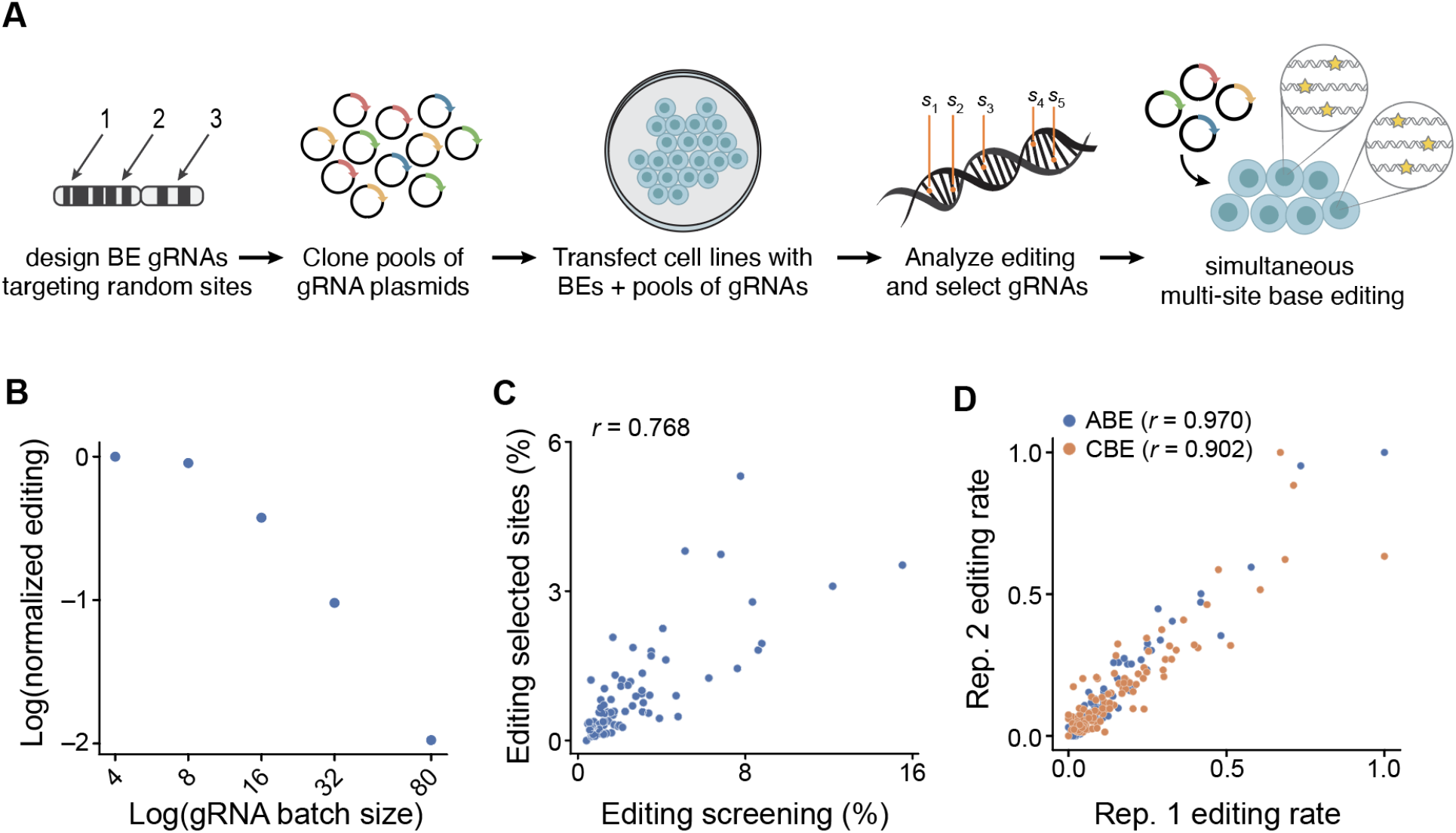
Strategy for simultaneous multi-site genome editing. **a.** Overview of design and evaluation of gRNAs workflow. Base editor gRNAs targeting random genomic sites were designed. Their editing efficiency was evaluated using pooled cloning and transfection. Editing rates were analyzed through sequencing, and gRNAs with high editing efficiency and low background were selected to be used for simultaneous multi-site base editing. **b.** Correlation of editing rate and transfected gRNA batch size. Editing rates were compared using the same gRNAs for different batch sizes of gRNA, with the total concentration of gRNAs per transfection kept constant. Editing efficiencies were normalized, and the decrease in editing rate per site (log-fold change) was calculated. **c.** Correlation between editing of gRNAs cloned and transfected in pools (x-axis) and subsequently selected gRNAs that were individually cloned, quantified, combined in a pool, and re-transfected (y-axis). **d.** Replicate correlation of editing rates between sites for adenine base editors (ABEs) and cytosine base editors (CBEs).

We first characterized how the number of distinct gRNA plasmids impacted the editing efficiency per site. We assessed pool sizes of up to 80 gRNAs and found that increasing the number of gRNAs led to a decrease in the relative editing efficiency per site (**Fig. 1b**), when the mass of total gRNA was kept constant. Except for the two smallest pool sizes (four and eight gRNAs), this relationship was observed for all pool sizes we tested. This result suggests that the number of edits that can be installed simultaneously at a given detection threshold is constrained by the gRNA pool size.

Next, we used pooled cloning and transfection of gRNAs to evaluate editing at endogenous genomic sites. In two independent experiments with varied gRNA pool compositions, we found editing rates of the pooled assay to be strongly correlated with those of individually purified gRNAs (r = 0.768; **Fig. 1c, Extended Data Fig. 1a**). This result indicates that pooled screening is an efficient strategy for rapidly evaluating gRNA efficiency. Notably, predictions made by existing models of gRNA editing efficiencies did not correlate with our experimental results (**Extended Data Fig. 1b**)^27^. This finding suggests that multi-site editing at endogenous loci is determined by factors not captured by current models and highlights the need for experimental evaluation. We correlated two replicates to investigate the robustness of the obtained editing efficiency values for pools of individually purified gRNAs. We found that pooled editing rates for both CBEs and ABEs are highly correlated between two biological replicates (r = 0.902 for CBE, r = 0.970 for ABE (**Fig. 1d**). These results demonstrate the reproducibility of a one-step method for installing multiple base edits.

### Reliable encoding and detection of edits across >100 genomic sites

The ability to encode information in a single editing step can only be achieved if edits can be robustly installed and reliably decoded. Messages—specific edited states across sites—are encoded in mammalian cell lines through pooled transfections using a subset of gRNAs. In the present study, we chose a binary encoding scheme wherein bits corresponding to zeros are unedited reference bases and bits corresponding to ones are C•Gs converted into T•As, or A•Ts converted into G•Cs, respectively (**Fig. 2a**). We reasoned that in this binary implementation, approximately half of all sites would be edited for each full-length message and that we could best analyze the fidelity of message encoding and decoding by transfecting two even-sized, non-overlapping gRNA pools.

**Figure 2.**
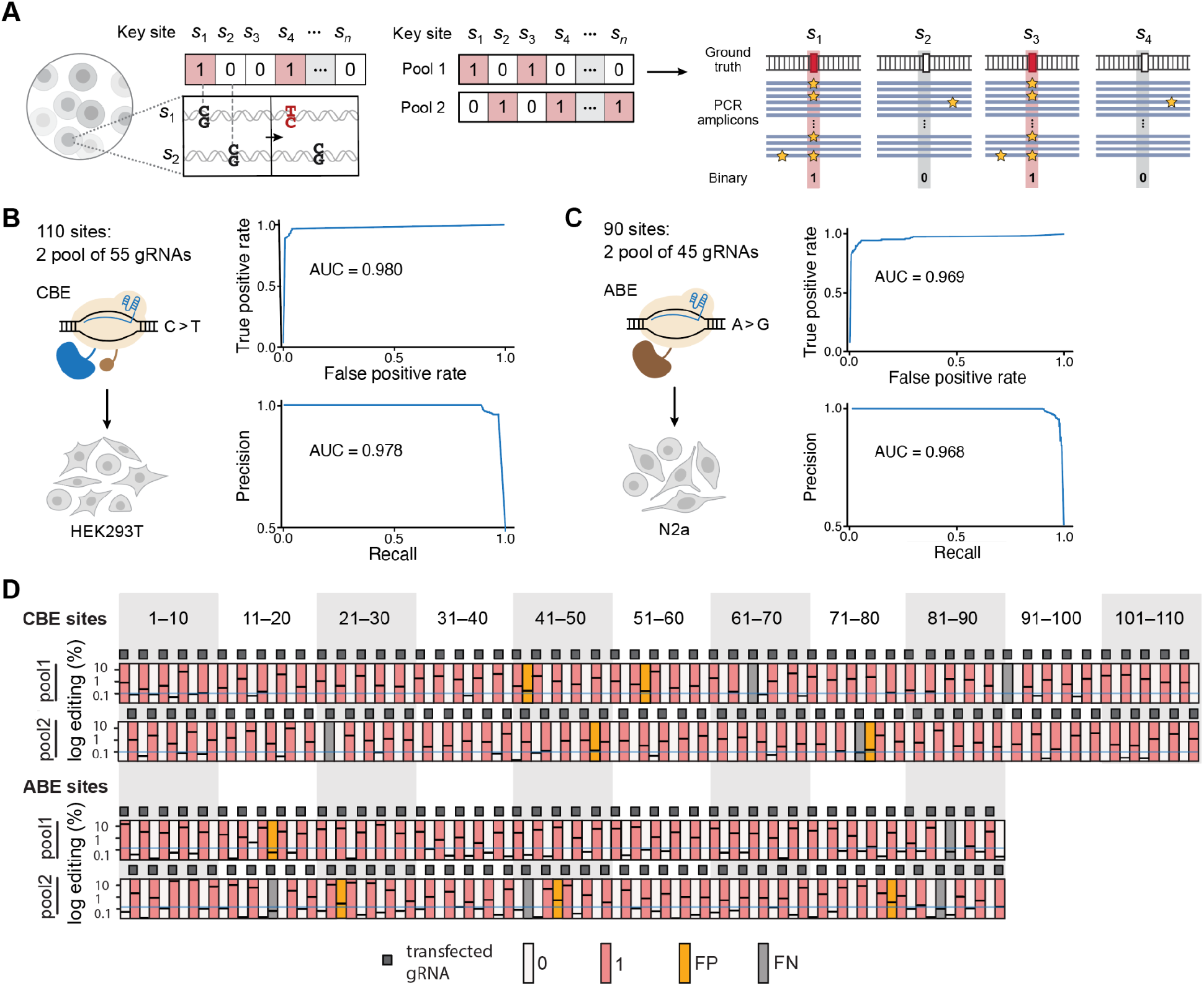
Multi-site base editing for information encoding. **a.** Left: Binary information encoding (here with CBEs) through edited or unedited state, corresponding to 1s and 0s, respectively. Right: gRNAs are split into two even-sized pools, transfected, and edits are detected through targeted high throughput sequencing **b,c.** ROC (top) and precision-recall (bottom) curves for detecting gRNAs transfected in pools of 55 and 45 gRNAs each for CBE and ABE, respectively. **d.** Classification of editing outcomes with selected editing threshold. The blue line represents the editing threshold (0.1% editing for CBE and 0.145% editing for ABE). Observed editing values are shown as black dashes. True positives and true negatives are shown in red and white, respectively, and false negatives (FNs) and false positives (FPs) are colored in grey and yellow, respectively.

To analyze the fidelity with which intended edits are installed and detected by an intended recipient, we determined whether we could correctly identify which gRNAs were transfected using targeted sequencing. We calculated receiver operator characteristic (ROC) curves for varying editing thresholds and defined true positives (TPs) as the transfected gRNAs that showed detectable editing and true negatives (TNs) as the non-transfected sites for which no editing was detected. Given that not all transfected gRNAs will yield editing efficiency values above an allele frequency threshold and that sites with no gRNA transfection might exhibit some background, we expect these cases to introduce classification errors (false negatives (FNs) and false positives (FPs), respectively). To determine CBE efficiency, we evaluated a total of 110 sites. We calculated the area under the ROC curve to be 0.980, indicating that targeted and untargeted sites can be classified with high accuracy (**Fig. 2b**). Precision-recall analysis further showed an area under the curve of 0.978, indicating that it is possible to detect edited sites with high accuracy (high precision) as well as the majority of positive results (high recall). To determine ABE editing efficiency, we evaluated a total of 90 gRNAs. We calculated the area under the ROC curve to be 0.969 and the area under the precision-recall curve to be 0.968 (**Fig. 2c**). These results demonstrate that one-pot encoding with CBE and ABE allows an intended recipient to discern edited and unedited sites with high accuracy.

To determine a threshold for detecting edited sites in a population of cells, we chose an editing rate that minimizes misclassifications based on data where 50% of sites were targeted. For CBE, we found that an editing threshold of 0.1% gave us optimal results with only 4/110 false negatives and 4/110 false positives, corresponding to a TP rate of 96.4% and a TN rate of 96.4% (**Fig. 2d**). For ABE, an editing threshold of 0.145% yielded best results 4/90 false negatives and 4/90 false positives, corresponding to a TP rate and TN rate of 95.6%, respectively. Our results demonstrate that simultaneous multi-site base editing provides a facile method for reliable information encoding and retrieval across >100 sites in the mammalian genome.

### Genomic Sequence Encryption (GSE) is a robust method for cryptographic security

To demonstrate that GSE provides information security, we computationally simulated the difficulty of breaking the encryption. The cryptographic key in GSE comprises the genomic indices at which mutations are installed, which we also refer to as ‘key sites’. Without access to the key sites, breaking the encryption (i.e. detecting the key sites) requires searching over the entire length of the genome. This process is prone to overlooking edited sites and detecting false hits as a result of sequencing errors, single nucleotide polymorphisms (SNPs), or artifacts that can occur during library preparation (**Fig. 3a**.). We first modeled the difficulty of this brute-force decryption for a single message, then over multiple messages to finally determine the cost of breaking the encryption.

**Figure 3.**
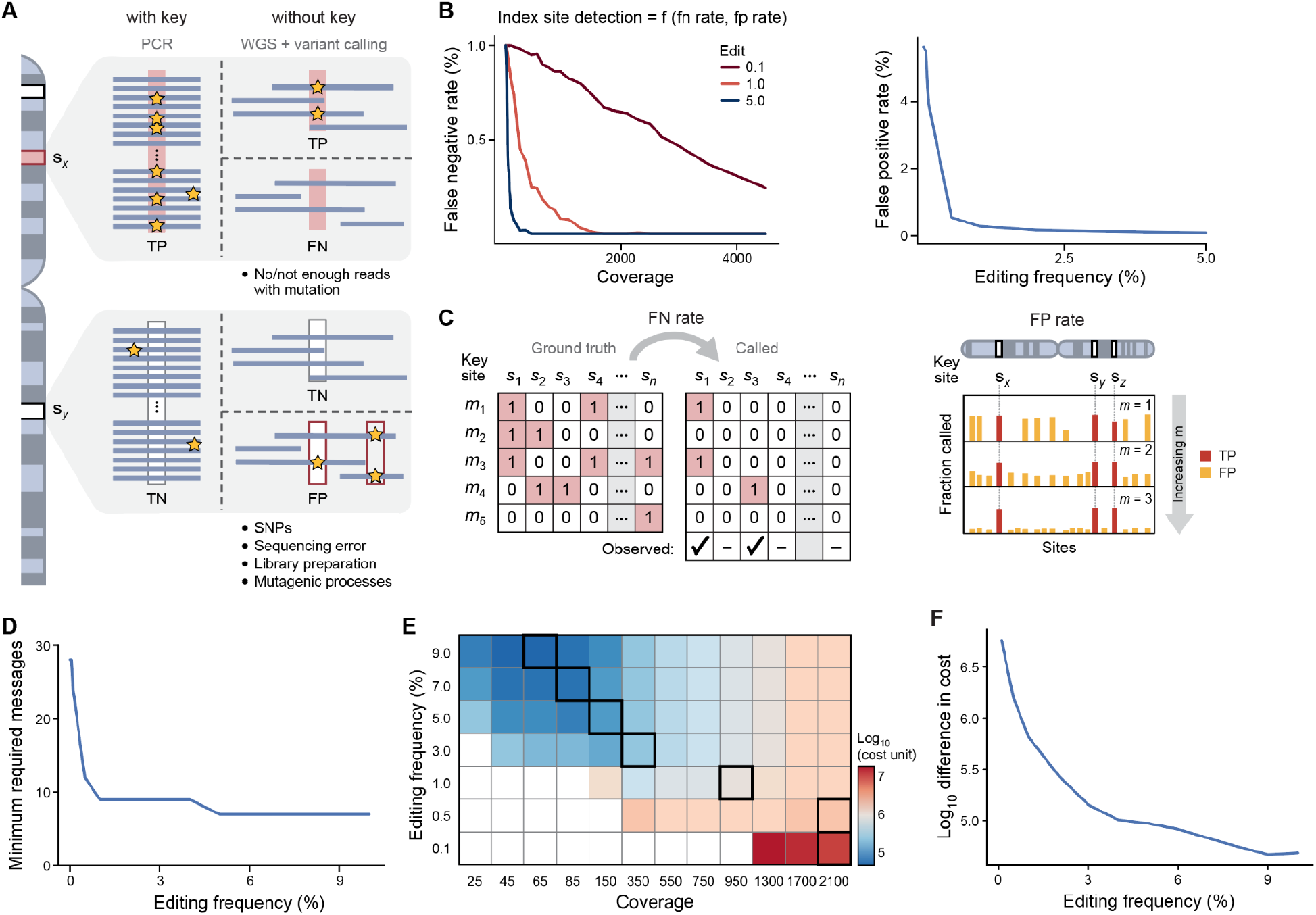
Secure information transfer through the asymmetric difficulty of detecting point mutations. **a.** When the key is unknown, whole genome sequencing (WGS) and variant calling software are required to detect installed edits. Possible outcomes are true positives (TPs), false negatives (FNs), false positives (FPs), and true negatives (TNs). **b.** The detection rate of an edited index site without a key depends on the false negative rate and false positive rate. The false negative rate depends on coverage and editing frequency, while the false positive rate depends on editing frequency. **c.** Breaking the key over multiple messages. Left: Detection of TPs is obscured through false negative rates, but sites can be detected over multiple messages. Right: Distinction between false positives and true positives over multiple messages, with TPs in red and FPs in yellow. **d.** Minimum number of messages required to break the key for an adversary when not limited by sequencing coverage. **e.** Sequencing cost to break the key over different editing frequencies and coverage levels (log_10_). The coverage level required for breaking the code within 30 messages for each editing frequency is boxed in black. **f.** The difference in cost in detecting the message without versus with key over various allele frequencies.

We developed a simulation framework in which we introduced synthetic edits into a published human deep sequencing data set and examined the performance of commonly used variant callers (**Methods**). We evaluated the detection of introduced mutations (false negative rate), the incorrect detection of variants (false positive rate), and their relation to the allele frequency of edits. The false negative rate was inversely proportional to both the allele frequency of the edit and sequencing coverage (**Fig. 3b**, **Extended Data Fig. 2a**). The false positive rate was dependent on the variant caller sensitivity and, thus, indirectly on the allele frequency of the edits (**Fig. 3b**), as the sensitivity would need to be adjusted to allow the detection of true edits. At a sensitivity of 0.1%, the false positive rate corresponded to ∼4% or millions of sites across the human genome. These results suggest that an unintended recipient would be unable to discern index sites by observing a single message. Next, we experimentally validated the difficulty of detecting edited sites at unknown coordinates by inducing and analyzing edits across the human exome at >1000x coverage. We observed both a decrease in performance in detecting edits (**Extended Data Fig. 2b**) and a simultaneous increase in false positive rates (**Extended Data Fig. 2c**) as allele frequencies decreased, indicating that detecting key index sites remains challenging even at a high level of coverage.

In an ideal cryptographic system, the same key can be used over multiple messages while still providing secure information transfer. We hypothesized that observation of multiple messages would enable us to distinguish TPs from FPs that are due to sequencing errors and library artifacts: While TPs are repeatedly detected over several messages, FPs are randomly distributed in the genome (**Fig. 3c**, **Extended Data Fig. 2d**). We simulated this scenario (under the assumption of infinite sequence coverage) and found that a minimum of 8 messages is required to break the code. As allele frequencies decrease, the number of messages needed to break the encryption scheme further increases. At an editing frequency of 0.1%, an unintended recipient would need to observe at least 24 messages (**Fig. 3d**) to break the code. GSE is, therefore, virtually impossible to break and enables secure information transfer until a certain number of messages is observed (the number is dependent on the allele frequency of the installed edits).

Finally, we investigated the feasibility of breaking the encryption if the number of messages observed is higher than the number required to provide complete security. We modeled this scenario under ideal conditions for the unintended recipient, assuming knowledge of editing allele frequencies, and calculated the total cost for two scenarios. In the first scenario, an unintended recipient can freely vary sequencing coverage and the number of observed messages to achieve the lowest cost (**Extended Data Fig. 3a**). In the second scenario, an unintended recipient needs to break the code within fewer than 30 messages (corresponding to a more realistic scenario in which a limited number of messages are observed, **Fig. 3e, Extended Data Fig. 3b**). Compared to the cost of a recipient who has access to the key (**Fig. 3f**, **Extended Data Fig. 3c**), the cost of breaking the encryption is ∼10^5^ higher at allele frequencies of 5%. This cost difference increases non-linearly at lower allele frequencies. At an editing frequency of 0.5% (well above the utilized detection threshold of 0.1%), the cost difference between recipients with and without the key is greater than 10^6^.

Taken together, these results show that the encrypted message remains largely inaccessible without prior access to the key. Revealing key indices requires the observation of multiple messages, after which it is asymmetrically difficult to break the code compared to someone with the cryptographic key. Finally, GSE adheres to Shannon’s maxim for secure cryptographic systems since the message remains secure even if the information encoding method is known.

### Encrypted cell signatures to detect instances of further genomic manipulation

We demonstrate an application of GSE for cell signatures—short messages in genetically modified strains that an intended recipient can easily read out to authenticate a cell strain. Additionally, we show that GSE enables editing at a ‘quality control’ (QC) site to determine whether a cell line has been genetically modified in the interim (**Fig. 4a**). Genetic modifications routinely involve transfection and selection steps that require/lead to a reduction in cell population size. This size reduction can lead to shifts in the genetic composition (genetic bottleneck) in the cell population, causing perturbations in the editing frequency at the QC site. For an unmodified strain, on the other hand, the allelic frequency would be expected to remain stable. We, therefore, reasoned that by creating an edit at the QC site with a defined allelic frequency (**Extended Data Fig. 4**), a recipient of a cell strain could detect whether a population of cells has been subjected to a genomic bottleneck. Such a capability is akin to computational cryptographic schemes, which frequently incorporate a method to validate the integrity of the data.

**Figure 4.**
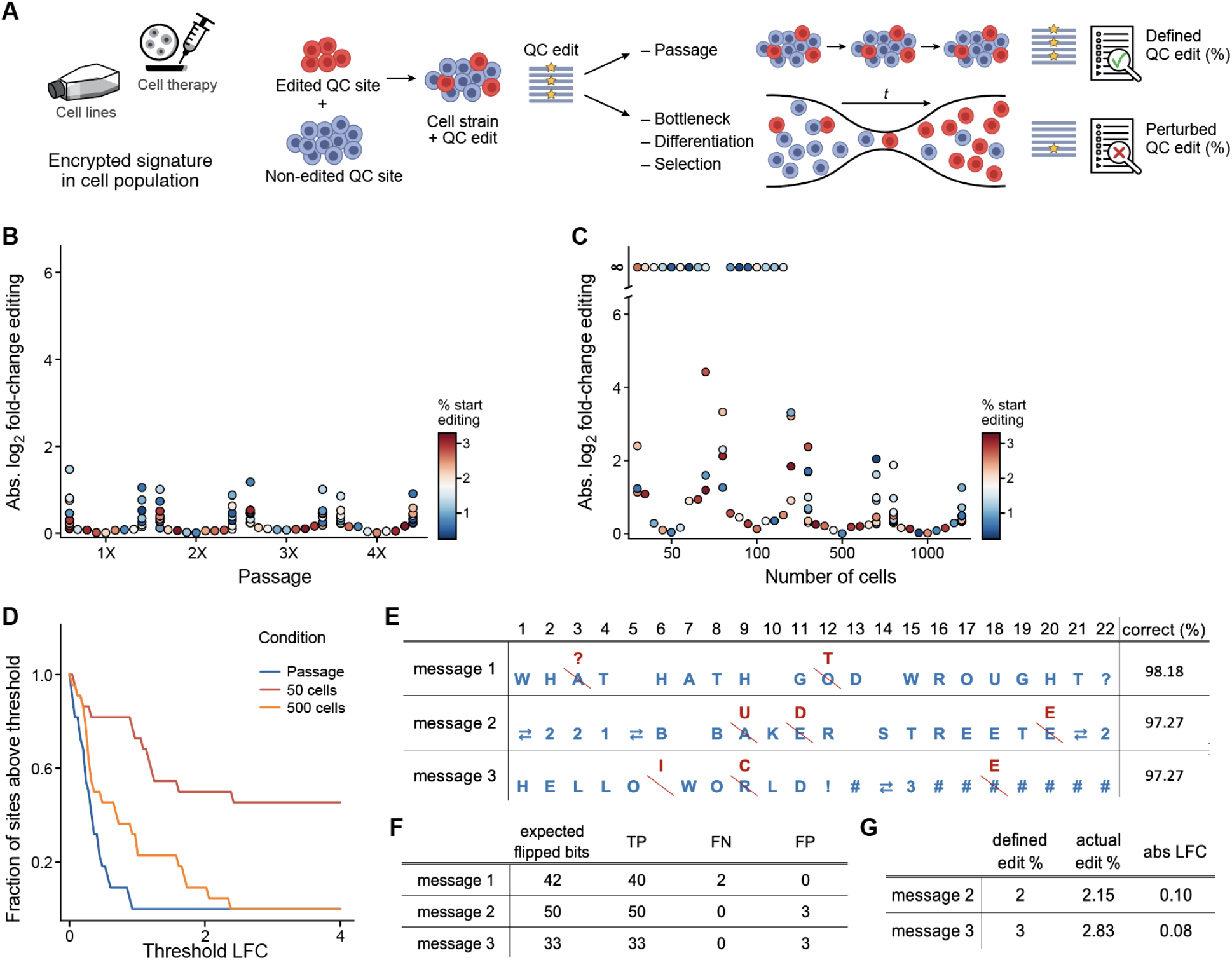
Message encoding and detection of bottlenecks through allelic frequencies. **a.** Overview of the application of GSE for encrypted cell signatures in cell populations (such as biotechnological cell lines or cell therapy). Under regular growth conditions, the editing frequency at the QC site is maintained. However, when cells are subjected to a bottleneck, the editing frequency is perturbed. **b.** Absolute log_2_ fold change (LFC) of editing frequency under regular passaging conditions. **c.** Absolute log_2_ fold change (LFC) of editing when bottlenecked to 50, 100, 500, or 1000 cells. **d.** Fraction of sites with editing changes above different log_2_ fold change (LFC) thresholds for cells after four passages and cells that were bottlenecked to 50 and 500 cells, respectively. **e.** The original message is shown in blue, and errors occurring during encoding or decoding are shown in red. (The double errors represent the shift character for shifting to numeric values.) **f.** Decoding of messages, showing the number of true positives (TP), false negatives (FN), and false positives (FP). **g.** Cell strains were mixed with a strain that carries edit at the QC site. ‘Defined edit (%)’ is the desired editing percentage as encoded in the message, and ‘actual edit (%)’ is the experimentally observed percentage.

First, we examined changes in editing frequency when a population of cells is subjected to a genetic bottleneck compared to regular growth conditions. We created a mammalian cell line with silent mutations and compared shifts in editing frequencies when cells were bottlenecked or maintained under regular conditions (**Fig. 4b**). We observed that edits remain more stable under regular passage conditions, where the highest absolute change in editing frequency was by a factor of 2.78 at any of the passages. In contrast, for the cell population bottlenecked at 500 cells, the highest absolute change observed was 5.18-fold. Subjecting cell populations to more stringent bottlenecks led to a larger disruption of editing frequencies (**Fig. 4c**). Additionally, we observed that 8/20 and 6/20 edited sites were no longer detected after the population was bottlenecked to 50 and 100 cells, respectively. We next calculated the fraction of sites for which the editing percentage was perturbed over different log fold change thresholds (**Fig. 4d**). We observed that the majority of sites in a 50-cell bottlenecked population were perturbed by greater than a 2-fold change in allele frequency. In contrast, when cells were passaged over 12 days, none of the sites had a greater than 2-fold change in allele frequency. These results illustrate that perturbations of editing rates depend on the maintenance conditions of a cell strain and that we can use allelic frequency at the QC site to determine whether a strain has been subjected to bottlenecks.

Next, we demonstrate that we can encode short messages and install an edit at the QC site at a desired editing frequency. We employed a modified version of the five-bit International Telegraph Alphabet no. 2 (ITA2) for converting text to binary. Three messages were selected for encoding, including the expected editing value at the QC site at the end of the message: ‘HELLO WORLD!#3’, ‘WHAT HATH GOD WROUGHT?’ and ‘221B BAKER STREET#2’. After transfection, an edit at the QC site was added at a defined editing percentage by mixing the cell population containing the encoded message with a cell population that contained an edit only at the QC site. For each of the three messages, less than 3% of all bits were misclassified (**Fig. 4e**). Editing values at the QC site were within ∼10% of the desired editing frequencies (**Fig. 4f)**, demonstrating that defined editing frequencies can be achieved.

These results demonstrate that messages can be successfully encoded and retrieved using pooled multi-site base editing, even when a naive encoding scheme with a uniform threshold for edit detection and no error correction mechanism is used. Furthermore, our results show that analyzing editing rates at the QC site can reveal whether a strain has been subjected to a bottleneck.

### Generation of individual embryonic stem cells (ESCs) with over two dozen edits

To extend encrypted signatures to living animals, multiple edits need to be made in the genome of a single cell. Given that mouse embryonic stem cells (mESCs) are routinely injected into zygotes to generate transgenic mice, we hypothesized that encoding a signature in mESCs would extend the application of encrypted signatures to living animals (**Fig. 5a**). As stem cells are known to be less amenable to genome editing than commonly used cell lines^28^, we investigated the feasibility of simultaneously editing multiple sites in mESCs. We installed a short genomic signature (“EUREKA”) by transfecting mESCs with base editor and a pool of thirteen gRNAs. After one round of editing in a cell population, we observed editing across all targeted sites, and an additional round of editing further increased editing rates by an average of ∼30% per site. Sites with low initial editing frequencies had increases in editing of up to ∼100%. Meanwhile, sites with higher initial editing frequencies showed more moderate increases in editing frequency (**Extended Data Fig. 5a**). These results demonstrate that it is possible to encode short genomic signatures in a population of mESCs and that editing rates can be increased by iterative editing.

**Figure 5.**
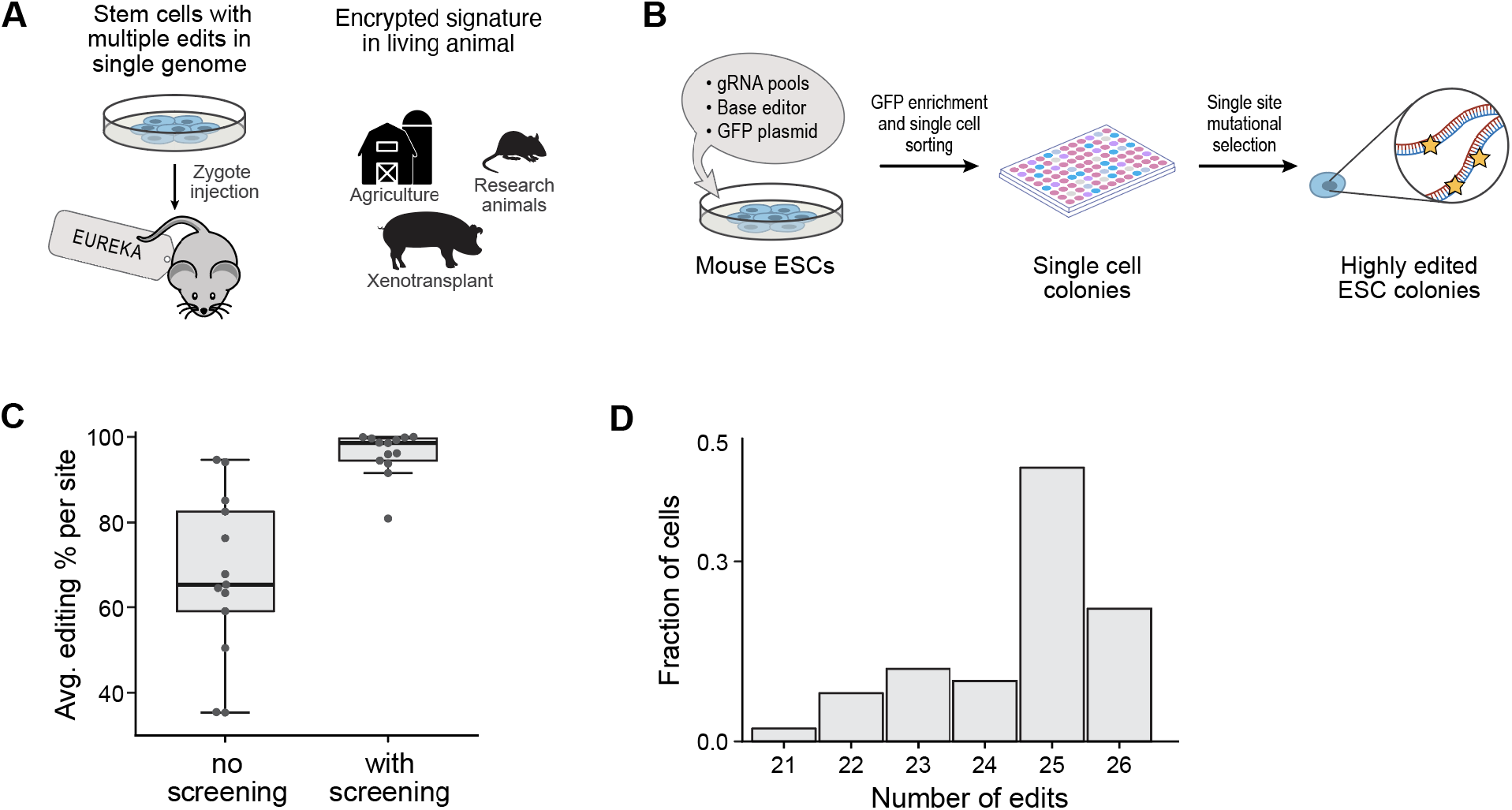
Multi-site editing in individual embryonic stem cells. **a.** Overview of the application of GSE for encrypted signatures in living animals. Obtaining cells with mutations within a single genome represents the first step toward extending GSE to living animals. mESCs are routinely used for zygote injection and can thus be used to generate transgenic animals; here we aim to install the message “Eureka” in mESCs. **b.** Overview of protocol for obtaining individual editing cells. **c.** Editing across all sites before enrichment and after enrichment. **d.** Histogram of the total number of edits per cell after enrichment for editing at a single site (out of a total of 26 edits).

Next, we sought to determine whether individual cells carrying a mutational signature could be obtained through an enrichment protocol (**Fig. 5b**). As population-level editing rates do not allow us to determine the zygosity of edits, we isolated and cultured individual cells from an edited cell population. As only one copy of the genome would be passed on via the germline, a single-cell colony with a full set of homozygous edits is needed to pass on all edits of the encrypted signature to the progeny. First, we investigated whether co-transfection with GFP can be used to enrich highly edited cells by selecting the top 5% GFP-expressing cells. We analyzed editing at the site with the lowest editing frequency and observed an increase in editing from 35% to 67%. This result suggests that enrichment of the top 5% GFP signal is an easy step to increase editing (**Extended Data Fig. 5b**).

To enrich for highly edited cells, we sought to determine whether co-editing between sites would allow us to obtain single cells containing all edits. We hypothesized that edits are not evenly distributed across individual cells, but that instead cells carrying one edit would be more likely to show editing at other genomic sites (due to factors such as transfection efficiency or the amount of expressed BE protein per cell). We opted to enrich for cells that are edited at the site that is most rarely edited in individual cells, i.e. the site with the lowest editing frequency in the cell population. As editing efficiencies vary between gRNAs, we reasoned that a cell that carries an edit at a low-efficiency site would be more likely to also carry edits at other genomic sites (with higher gRNA editing efficiencies). Selecting clones that carry an edit at the site with the lowest editing rate, resulted in highly enriched editing and increased the average editing rate across all sites from 67 to 96% editing (allelic frequency of edits, averaged across all sites) (**Fig. 5c**).

We next analyzed the zygosity of single-cell clones that carry an edit at the site with the lowest editing rate. 100% of the analyzed clones (50/50 clones) had editing across all of the targeted sites; We detected either heterozygous or homozygous editing outcomes and did not detect any unedited sites among the enriched single-cell clones. The analyzed cells carried an average of 24.5 out of 26 edits (13 sites x 2 in a diploid genome) (**Fig. 5d**). The most frequent number of edits per cell was 25 edits (in 46% of cells). The second most common number of edits was 26 edits (in 22% of cells), and the lowest observed number of edits per cell was 21. These results suggest that screening for editing at the site with the lowest editing frequency allows for easy identification of multi-site edited clones; cells with homozygous edits at all targeted sites can be identified with minimal screening. Collectively, these results further demonstrate that it is possible to isolate individual cells that are highly edited with minimal screening. Identifying a site with low editing efficiency compared to other targeted sites enables a facile screening method for highly edited cells. Furthermore, our results demonstrate that signatures can be installed in individual mESCs.

## DISCUSSION

In this study, we introduce GSE, the first DNA sequencing-based cryptographic system. We demonstrate that gRNA pools enable robust multi-site editing and implement GSE using base editing across >100 genomic sites in mammalian cells. With a cryptographic key (knowledge of the genomic coordinates), a recipient can decode messages through targeted sequencing. Our computational simulation and experimental data illustrate that decoding without the key is impossible until a certain number of messages are observed. Beyond this threshold, breaking the key is theoretically feasible but comes at a significantly higher sequencing cost; this asymmetric sequencing cost resembles current computational encryption algorithms. We outline an application of GSE for the authentication of a cell line/biological organism by reading its falsification-proof genomic signature. Additionally, we demonstrated that allelic frequencies can be leveraged in a cell strain to verify the absence of genomic alterations in the interim, akin to digital authentication of message integrity. Therefore, this work represents a completely biological instantiation of many key concepts in modern cryptography in mammalian cells. Finally, we developed a protocol for enriching highly edited individual stem cells and demonstrated that we can obtain individual mESCs with >25 genomic This capability extends the applicability of encrypted signatures to living animals.

We show that base editors are suitable for introducing multiple simultaneous edits in mammalian cells. Base editors have been suggested to enable a higher number of edits within a single genome, overcoming the toxicity barrier of traditional Cas9^13^. Previous efforts to introduce multiple edits in primary and stem cells have been limited to no more than five distinct edits simultaneously^25,26^. In this study, we demonstrate that we can obtain stem cells with 26 edits across a single diploid genome, by selecting cells that are edited at the site with the lowest editing efficiency. This advancement anticipates applications in biotechnology and fundamental biology, including probing multi-site genomic interactions at single-base resolution or the gene level^13^. While we have yet to test this step for other gRNA pool sizes and cell types, we anticipate that our protocol would enable the selection of cells bearing a higher number of mutations and may be generalizable to other cell types. Additionally, coupling the low-efficiency edit with a selectable phenotype, e.g., an antibiotic or fluorescent marker, will further facilitate the selection of highly edited clones.

We demonstrated the encoding of up to 110-bit messages through base editing. We anticipate that longer messages can be encoded simultaneously by screening for a larger number of gRNAs with high editing efficiencies. Editing efficiency per site decreases with an increasing number of gRNAs in the pools; there are limits to the number of edits that can be made within a single round. This challenge can be circumvented through iterative rounds of editing. While the base editors used in this study require the presence of an NGG PAM site – thus imposing some restrictions on targeted sites – newer evolved base editors with relaxed or altered PAM site requirements^29,30^ could be employed to expand the accessible sequence space. More recently, the development of novel base editors that enable base transversions^31–33^ allows for the introduction of nearly all types of point mutations. In addition, we expect that the scheme could be expanded to prime editors^34^.

GSE relies on orthogonal theoretical underpinnings to existing cryptographic approaches and does not rely on computational difficulty. It is therefore not affected by increases in computing performance as well as the emergence of quantum computing, and decryption via Shor’s algorithm. Additionally, GSE can be extended to include other information security concepts, such as ‘winnowing and chaffing’, in which additional edits that do not include information are installed to add noise^35^. A sparser encoding, where a lower fraction of the total sites is edited, would further increase the difficulty of breaking the encryption. Moreover, we envision that by targeting sites of common genetic variation, the message would become practically indistinguishable from SNPs.

GSE presents a versatile encryption scheme that can be generalized to other organisms amenable to multi-site genome editing. As genome engineering and genetically modified organisms (including cell lines, animals, and crops) become more widespread, we anticipate that approaches like GSE for securing and authenticating biological resources will become increasingly important. Additionally, we expect that our work on multi-site editing including the selection of highly edited cells will be of relevance for various applications in biotechnology and fundamental biology.

## Acknowledgments

We thank Luyi Tian, Dylan Cable, Seth Shipman, Kai Liu, Vijay Sankaran, Jorge Martin-Rufino, Hattie Chung, Siddharth Iyer, Zijay Tang, and members of Chen and Church labs for helpful discussions.

## Author contributions

V.V. and F.C. conceptualized the idea. F.C. and G.M.C. supervised the study. V.V., S.Z., K.S. and S.Q. performed methodology, investigation and visualization for the study. V.V., S.Z., F.C., G.M.C. wrote the manuscript.

## Data and Material Availability

All sequence data will be deposited on sequence read archive upon publication. All plasmids will be made available on Addgene. Code used will be deposited to Github and Zenodo upon publication.

## List of Supplementary Materials

Supplementary Table 1: gRNA target sequences and sequencing primers.

Supplementary Table 2: Modified version of five-bit International Telegraph Alphabet no. 2 (ITA2) for converting text to binary.

## METHODS

### gRNA Design

For designing gRNAs, random genome indices were retrieved using bedtools (bedtools version 2.27.1) running the command ‘bedtools random’ on the human reference genome hg38. Corresponding fasta sequences were extracted and a custom Python script was used to design gRNAs as follows: The nucleotide sequence and its reverse complement were queried for 23 nucleotide sequences that have base C at positions 4-8 and bases NGG at positions 21-23 where N can be any of the four bases, corresponding to the PAM site requirement for SpCas9 which of AncBE4max. Sequences were further filtered to exclude guides with homopolymer stretches of four or more nucleotides and a G/C content of lower than 30%. Only one gRNA per site was selected.

### Cloning of gRNA Pools for gRNA Selection

For pooled cloning of gRNAs, gRNA sequence and adjacent bases were ordered as eblocks from IDT, and pools of 32 gRNAs each were cloned into the backbone pSB700 mCherry (addgene #64046) using Gibson cloning. Plasmid pools were then transformed into 5-alpha competent *E. coli* (New England Biolabs), and the *E. coli* culture carrying the gRNA plasmids was cultivated for ∼12 hours in LB medium before plasmids were extracted via mini-prep (New England Biolabs).

### Cloning of Individual gRNAs for Selected Sites

For selected sites, eblocks were cloned into the backbone pSB700 mCherry (addgene #64046) and transformed into 5-alpha competent *E. coli* (New England Biolabs). Cloning was performed for eight gRNAs at a time, and transformed *E. coli* cultures were subsequently plated on LB agar. Colonies with correct gRNA inserts were identified using Sanger sequencing and colonies with sequence-verified gRNA plasmids were subsequently individually mini-prepped for transfection.

### HEK293T and N2a Cell Culture and Transfection

HEK 293T cells and Neuro2a (N2a) cells were obtained from ATCC and were authenticated and tested negative for mycoplasma by the manufacturer. Cells were maintained in Dulbecco’s modified Eagle’s medium with Glutamax and Sodium pyruvate (Gibco) with 10% Fetal Bovine Serum (Gibco) at 37°C and 5% CO_2_. Medium was exchanged every 3 days and cells were regularly passaged before reaching ∼80% confluency using TrypLE (Gibco) for dissociation. For transfection, cells were seeded in 12-well plates 24 h prior, and transfected with Lipofectamine2000 (Thermofisher) according to the manufacturer’s instructions with modifications as outlined below: Cells in each well were transfected with 3 µg base editor DNA and 1 µg of gRNAs and using 5 µL of lipofectamine reagent. When multiple gRNAs were used in one transfection, gRNAs were pooled at equimolar concentrations. After transfection, cells were cultivated for 3 days and washed once with PBS before harvest.

### Targeted Sequencing and Analysis

Genomic DNA was extracted using a Zymo DNA extraction kit, and target sites were amplified in separate 25 µL PCR reactions using Kapa HiFi Hotstart readymix according to the manufacturer’s instructions with 125 ng of genomic DNA as a template.

Libraries were prepared using the NEBNextUltra library prep kit using a size selection step, pooled at equimolar concentration, and sequenced on an Illumina MiSeq, using paired-end sequencing.

Paired-end read fastqs were aligned to the human reference genome hg38 using bowtie2 version 2.3.4.3. The resulting aligned files were analyzed using a custom Python script. Base pileup for genomic indices corresponding to the key indices was performed using Pysam version 0.18.0, with minimum base quality set to 30. The fraction of edited bases was obtained by dividing the number of edited bases at the index position, i.e. T or As, by the sum of both reference bases, i.e. Cs or Gs, and edited bases.

### Sensitivity/Specificity Experiment: Decoding with Cryptographic Key

gRNAs were split into two evenly sized, non-overlapping pools by numbering gRNAs and splitting them into two pools of even-numbered and odd-numbered gRNAs, respectively. For CBE, 110 gRNAs were split into two batches of 55 gRNAs each, and HEK 293Ts were transfected with AncBE4max and gRNA batches as described above. For ABE, 90 gRNAs were split into two pools of 45 gRNAs each, and N2As were transfected with either of the two gRNA pools and Abe8e as described above. Edited sites were analyzed using amplicon sequencing and the analysis pipeline described above. The false positive rate at each editing percentage was calculated as follows: False positive rate = FP/(FP+TN). False negative rate was calculated using the following formula: False negative rate = FN/(FN+TP)

### Evaluating False Negative Rates

High-coverage sequencing data was downloaded from the SRA database SRX5342252. Fastq files were aligned to the human reference genome hg38 using bowtie2 version 2.4.1. Paired-end reads of the bam file were unpaired using a custom Python script and treated as single-end reads. Single base mutations were inserted using biostar404363 ([Lindenbaum, 2015](https://figshare.com/articles/journal_contribution/JVarkit_java_based_utilities_for_Bioinformatics/1425030/1)), at a distance of at least 450 bases to other artificial mutations to ensure independence of variant calling decisions. Allele percentages of synthetic mutations ranged from 0.001% to 20% at sites with sequencing coverage from ranges 10x to 5000x. 300 sites were chosen for each coverage level and one modified bam file was generated for each allele percent and coverage combination. Variant calling was performed and the false negative rate was determined, comparing two variant callers; Mutect2 and Varscan2. Varscan2 was run in somatic mode with the unmodified bam file as a normal control. The sensitivity flag ‘–min-var-freq’ was set to the allele frequency of the mutation and a minimum coverage level of 5x was required to call a variant.

### Evaluating False Positive Rates

We defined false positive cases as bases in the original unmodified bam that were called a variant with an allele percent lower than 30% (to exclude SNPs) and a required sequencing depth of at least five reads. The false positive count was divided by the number of bases with the required sequencing depth to derive the false positive rate. Varscan2 was run with pileup2snp and the sensitivity flag min-var-freq was set to a range of thresholds to derive the relationship between false positive rate and variant caller sensitivity; Mutect2 does not have a sensitivity flag.

### Modeling Difficulty of Breaking the Code without Cryptographic Key

While VarScan2 and Mutect2 had comparable performances at high allele frequencies, only VarScan2 was able to successfully detect variants at allele frequencies below ∼2% (**Fig. S3)** and we, thus, decided to focus the cost comparison on VarScan2 (Extended Data).

To evaluate the impact of the false negative rate on detecting mutations, we assume that a message converted into binary is randomly distributed as half zeros and half ones, and define the reveal rate as *Reveal Rate*(*RR*) = (1 − *FN*)/2, where FN is the false negative rate. We assume that if the adversary can discover 90% (threshold *T*) of the key indices, the key is considered broken. We calculate the number of messages (*m*) that an adversary has to observe to break the key: *m* = *log*(1 − *T*) / *log*(1 − *RR*). The final cost is calculated by multiplying the number of required messages times the cost for sequencing at the required coverage:

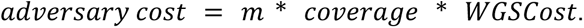

Next, we model the impact of the false positive rate. The variant caller false positive rate depends on its sensitivity setting which the adversary controls. We assume the adversary will have to set the sensitivity to at least be able to detect the lowest allele percent base edits. Experimentally the editing range is around 0.1% to 5%.

For each base depending on if it is a key index or not, the number of times it is called a variant over m messages will differ. Bases that aren’t key indices would be called a variant at the false positive rate while key index bases will be called at a rate equal to the reveal rate. These form two distinct binomial distributions with their mean at the proportion of times a base would be called a variant and sample size being the number of messages sequenced. The decision of whether a base is a key index or not is a question of which distribution it falls under. Key indices have their mean located at the reveal rate and non-key indices have their mean at the false positive rate. The overlap between the two distributions decreases with messages sequenced, and we set a threshold for the allowed overlap between the distributions that would allow the adversary to successfully uncover enough key indices to read or tamper with the message: The number of false positives an adversary could allow should be smaller or equal to the number of bits in the message, i.e. ∼100/3*10^9^. The threshold for allowable false negatives was set to 0.1, equivalent to the crack threshold defined above. A statistical pipeline was developed in RStudio to calculate how many messages the adversary needs to sequence to distinguish false positives from true key indices.

### Comparing Cost between Recipient with and without Key

We assume a cost of $1000 to perform whole genome sequencing (WGS) on the human genome with 30x coverage, corresponding to ∼$33 for WGS at 1x coverage. We calculate recipient cost as follows:

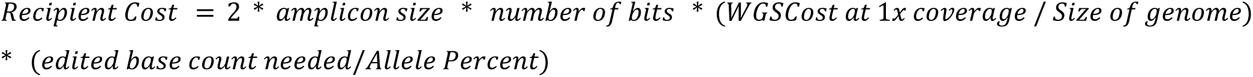

The estimated cost of sequencing a single base (WGS Cost at 1x coverage / genome size) is multiplied by the number of total bases needed to be sequenced to decide whether a key index encodes a 1 or 0. The total cost is doubled for paired-end sequencing. To determine the coverage needed to observe an edited site at least some number of times we divide that number by the allele percent.

### Evaluation of Stability of Editing Allele Frequency during Bottleneck

HEK 293T cells were transfected with gRNAs introducing silent mutations in the exome in batches of 4 gRNAs per transfection, and equal numbers of cells per transfection were pooled 3 days after transfection. For assessing stability over time, cells were maintained in a 12-well plate and passaged at a ratio of 1:8 every three days. At each passage, parts of the cells were harvested for genomic DNA extraction. For the bottleneck experiments, 50, 100, 500, or 1000 cells were sorted into 12-well plates at day 3 after transfection using a SONY SH800 cell sorter. Cells were cultivated for 14 days and harvested for gRNA extraction.

### Whole Exome Sequencing Library Preparation

Genomic DNA was extracted using a Zymo DNA extraction kit. A total of 500 ng of genomic DNA was fragmented using NEBNext Ultra II FS DNA Fragmentation module (New England Biolabs) for 20 min and 37°C, and whole genome library preparation was carried out using NEBNext Ultra II DNA Library Prep Kit, with a final PCR amplification step of 13 cycles. Subsequently, exonic regions were enriched using NextGen hybridization capture IDT using 500 ng library DNA as input and following the manufacturer’s instructions.

### Variant calling (Experimental Data) GATK Pre-Processing

Fastq files were aligned to the human reference genome hg38 using bowtie2 version 2.3.4.3. Next, bam files were processed following GATK4 best practices. Briefly, after alignment duplicates were removed and base quality scores were recalibrated using Picard. Processed bam files were then used as input for Mutect2 and VarScan2 and variants were called using a sensitivity threshold of 1% and 0.5% for VarScan2.

For determining the number of false positives, called variants with over 25% editing were filtered out, since those were assumed to be cell-line specific SNPs. The false positive rate was determined by dividing the number of false positive calls by the number of bases covered by the IDT hybridization panel.

### Mouse Embryonic Stem cell (mESC) cell culture and nucleofection

The mESCs used in this were ES-R1 (Sigma-Aldrich, #07072001). Cells were maintained in 2i+LIF media ([Sim, 2017](https://www.sciencedirect.com/science/article/pii/S2213671117301340?via%3Dihub)) on gelatine-coated plates at 37°C and 5% CO_2_. Plates were coated by incubation with 0.1% gelatine for an hour at 37°C and cells were passaged using Tryp(LE) Gibco for dissociation.

For nucleofection, cells were seeded 24 h before the experiment, washed once with PBS and 5*10^5 cells were nucleofected on the X-unit of a Lonza 4D nucleofector, following the manufacturer’s instructions, and using P3 Lonza nucleofection solution, one aliquot of dABE8e ([Richter,2020](https://www.nature.com/articles/s41587-020-0453-z)) and modified sgRNAs (Synthego) per well. gRNAs carried the manufacturer’s standard modifications (2’-O-methyl analogs (OMe) on the first and last three bases and 3’ phosphorothioate internucleotide linkages (PS) between the first three and last two bases), and RNAs were pooled at equimolar concentrations, to a total of 600 pmol per reaction. After nucleofection and recovery, cells were further cultivated in 2i+LIF on gelatine-coated plates for five days and washed once with PBS before sequence analysis. To assess iterative editing, cells were nucleofected seven days after the first nucleofection.

### Analysis and enrichment of single cells (mESCs)

For sorting, 1 ug of GFP was added to the nucleofection reaction mixture. Following nucleofection, mESCs were maintained for five days before cells were sorted (on a SONY SH800 sorter) into gelatine-coated 96-well plates, selecting the top 5% GFP-expressing cells. Cells were subsequently maintained in 2i media + P/S. Cells were maintained under regular media exchange until reaching ∼90% confluency when editing was analyzed by extracting genomic DNA using QuickExtract (Lucigen).

Editing at the site with the lowest editing frequency was analyzed, and single-cell clones with editing at this site were further expanded and their genomic DNA was harvested to analyze the other targeted sites. For analysis of editing rates per cell, only cells with sequencing reads for all of the site were analyzed; editing rates >95% were counted as homozygous edits and were otherwise counted as heterozygous when editing was <95% and >35%.

