## Extended Data Figures for "Cryptography in the DNA of living cells enabled by multi-site base editing"

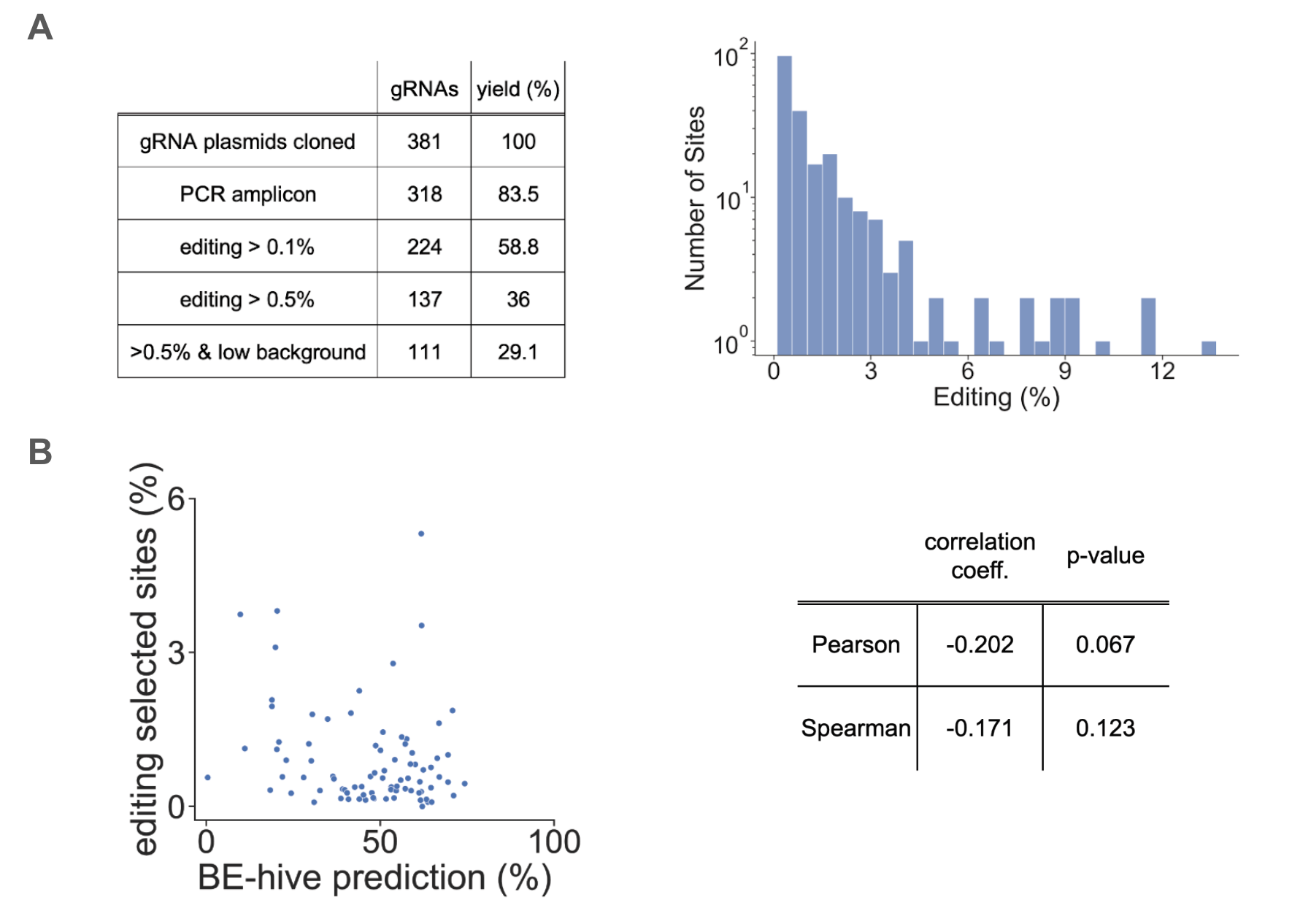

**Extended Data Figure 1.** Results of pooled gRNA cloning and screening for CBEs. **a.** Left side: Out of 381 gRNAs that were cloned, efficient PCR amplification of 318 of the corresponding genomic sites was achieved. When transfected in batches of 48 gRNAs, 224 sites showed editing greater than 0.1%, and 137 sites showed editing greater than 0.5%. We further analyzed the background rates of the 138 sites with > 0.5 editing (data not shown) and selected 111 sites with low background rates. Right side: Editing distribution of 226 sites that showed > 0.1 % editing when transfected in batches of 48 gRNAs, before filtering out sites with high background. **b.** Correlation between editing rates for BE-hive prediction (assuming editing with a single gRNA for BE4 in HEK293Ts) and observed editing rates in pools for selected sites. Pearson and Spearman correlation coefficients were calculated.

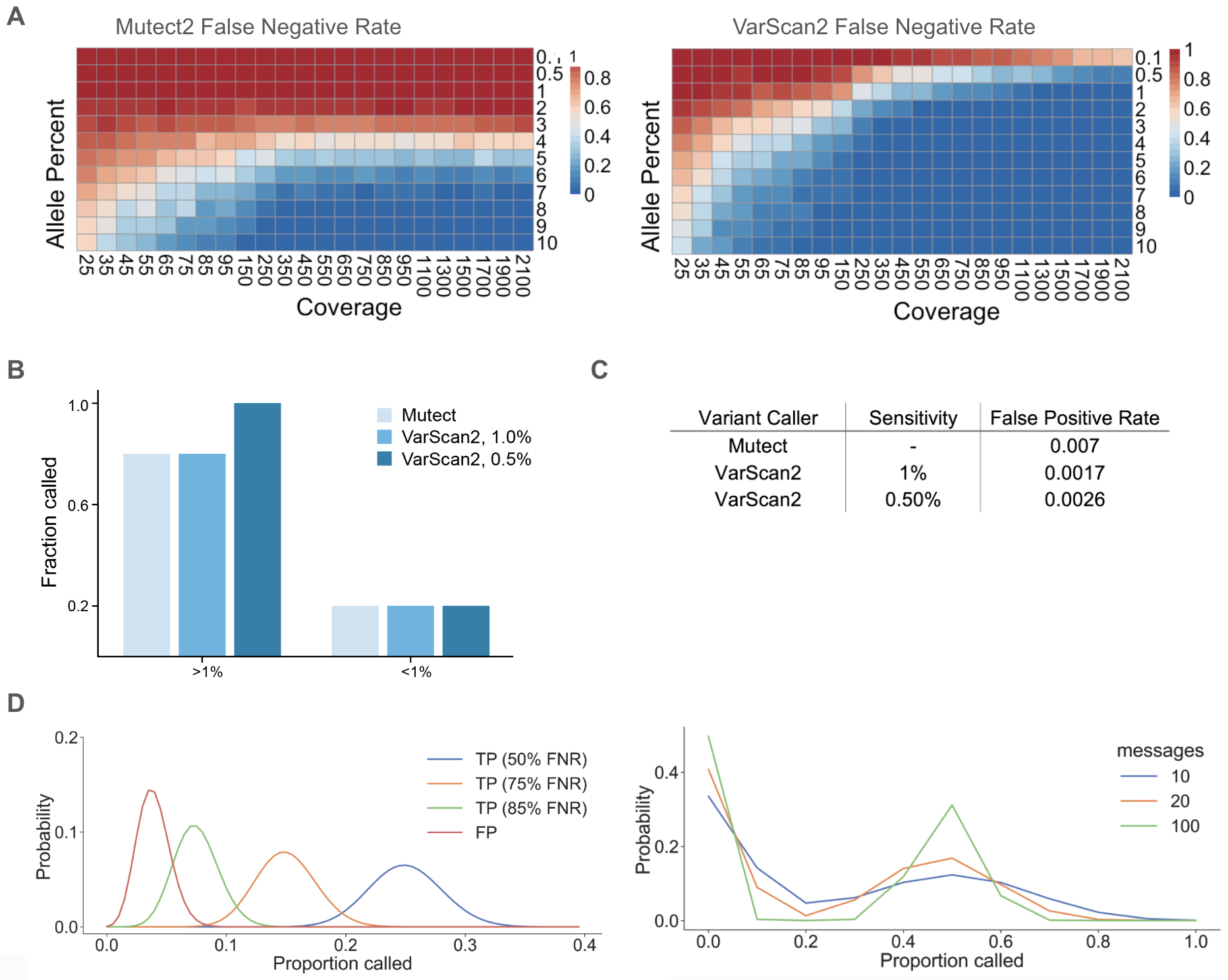

**Extended Data Figure 2.** Computational simulation and experimental results for the false negative rate and false positive rate (single message). a. False negative rates at varying allele percentages and coverages obtained from calling variants on read files with artificially introduced mutations. A. False negative rates obtained for Mutect2. B. False negative rates obtained for VarScan2. **b.** Fraction of detected mutations from whole exome sequencing data at ~1000x coverage with variant caller Mutect2 and VarScan2, at ~1000x sequencing coverage. **c.** False positive rates generated from whole exome sequencing. Whole exome sequencing was performed at 1000x coverage and variants were called using Mutect and VarScan at 1% and 0.5% allele frequency thresholds. **d.** The proportion of times a genomic index that is a true positive versus a false positive is called. Left: Binomial distributions of the number of times key indices would be called variants over 200 messages at false negative rates (FNR) of 50, 75 and 85%. Right: Binomial distributions of true positives and false positives and the proportion of times they are called variants over 10, 20, and 100 messages.

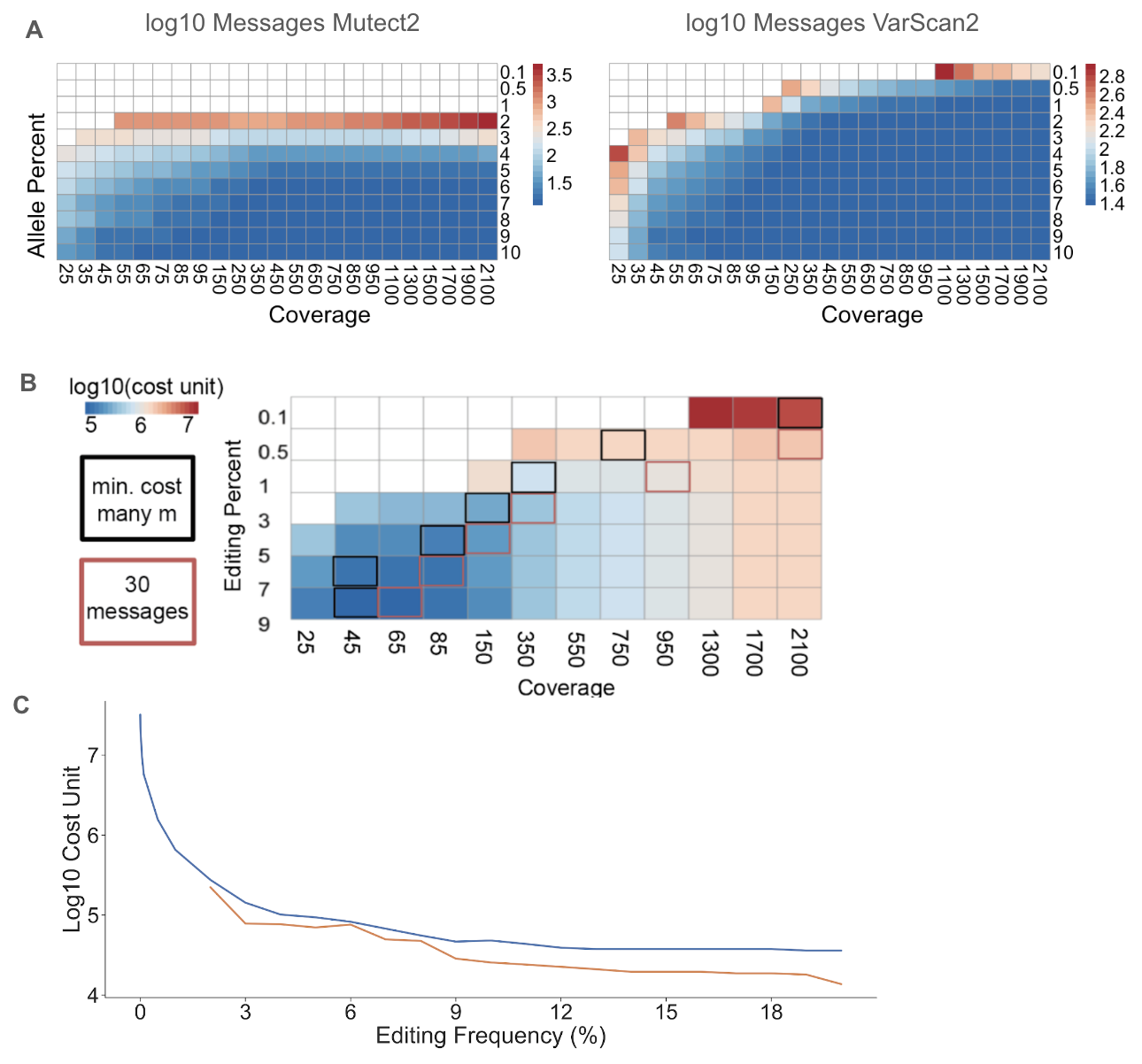

**Extended Data Figure 3.** Difficulty of breaking encryption scheme over multiple messages. **a.** Number of messages needed to reveal key indices using Varscan2 versus Mutect2. Values were calculated using a statistical analysis of how many messages are needed to meet an acceptable false positive rate of 100/(3*10^9) representing roughly the number of bits in a message over the size of the human genome and a false negative threshold of 0.1 meaning 90% of key indices have been discovered. **b.** Attackers cost when the optimal combination of sequence coverage and number of messages is chosen. Cost with unlimited messages is shown in black boxes; cost with messages limited to max. 30 messages is shown in red boxes. **c.** Cost for an adversary to break a key when messages are encrypted at various allele frequencies. The cost for VarScan2 is shown in blue and the cost for Mutect2 is shown in orange; at less than ~2% allele frequency the cost approaches infinity when using Mutect2.

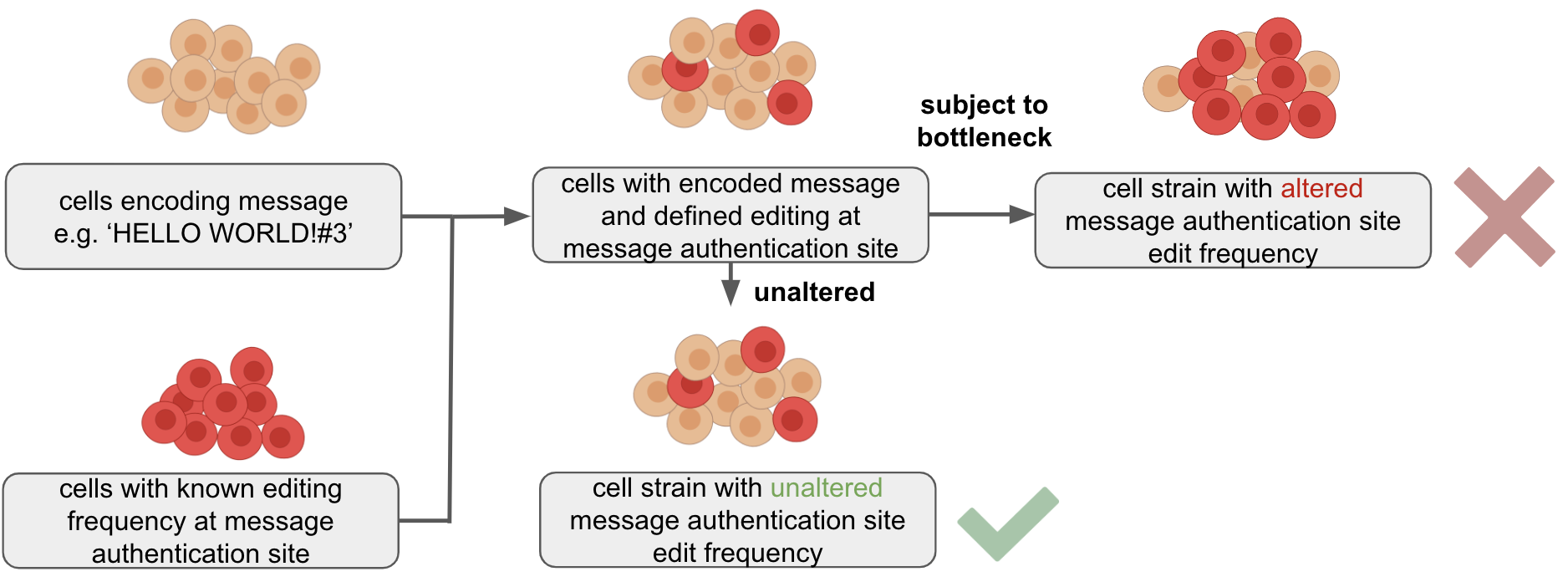

**Extended Data Figure 4.** Message authentication scheme. One of the key sites is specified as a message authentication site. A strain carrying a mutation at only this site is created, the editing frequency determined by sequencing, and the strain is then diluted into cell strains with encoded messages at a ratio achieving final editing frequencies as specified in the messages, e.g. in ‘HELLO WORD!#3’ the digit 3 specifies the editing frequency. Due to genetic drift, the editing frequency is expected to be perturbed when cells are subjected to a bottleneck compared to regular growth conditions, and the editing frequency can therefore inform about the integrity of the strain.

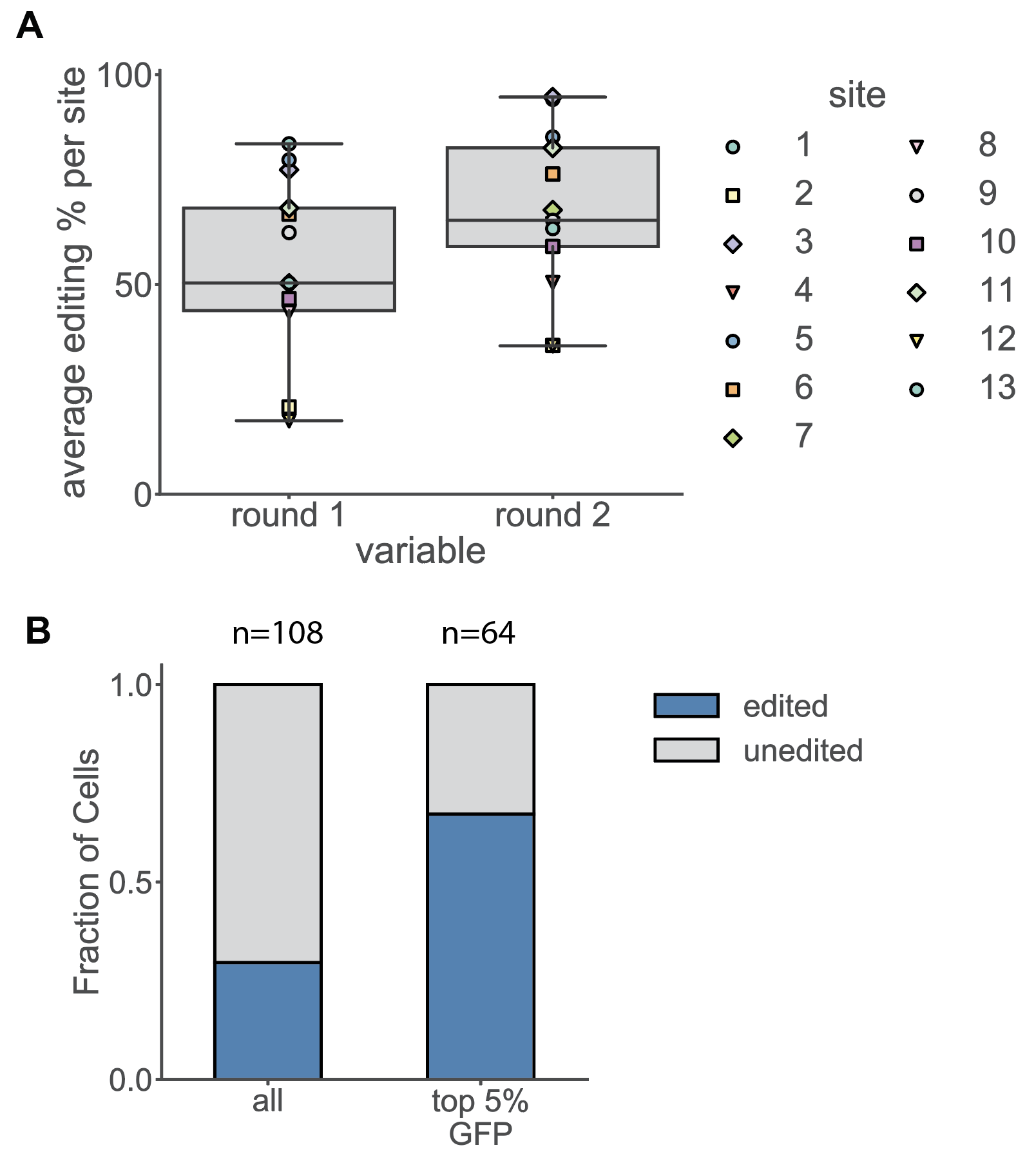

**Extended Data Figure 5.** Multi-site base editing in mESCs. a. Increase in editing per site over two rounds of editing in a population of mESCs. Each point represents editing frequencies for a single genomic locus, across a population of cells. **b.** Editing increase at a single genomic locus (the site with lowest editing efficiency) in top 5% GFP expressing cells, calculated from editing in single cell clones. n represents the number of cells that were sequenced for each condition.
